# An efficient context-aware approach for whole slide image classification

**DOI:** 10.1101/2023.01.15.524098

**Authors:** Hongru Shen, Jianghua Wu, Xilin Shen, Jiani Hu, Jilei Liu, Qiang Zhang, Yan Sun, Kexin Chen, Xiangchun Li

## Abstract

Computational pathology for gigapixel whole slide images (WSIs) at slide-level is helpful in disease diagnosis and remains challenging. We propose a context-aware approach termed <u>W</u>SI <u>I</u>nspection via Transformer (WIT) for slide-level classification via holistically modeling dependencies among patches on the WSI. WIT automatically learns feature representation of WSI by aggregating features of all image patches. We evaluate classification performance of WIT along with state-of-the-art baseline method. WIT achieved an accuracy of 82.1% (95% CI, 80.7% - 83.3%) in the detection of 32 cancer types on the TCGA dataset, 0.918 (0.910 - 0.925) in diagnosis of cancer on the CPTAC dataset and 0.882 (0.87 - 0.890) in the diagnosis of prostate cancer from needle biopsy slide, outperforming the baseline by 31.6%, 5.4% and 9.3%, respectively. WIT can pinpoint the WSI regions that are most influential for its decision. WIT represents a new paradigm for computational pathology, facilitating the development of effective tools for digital pathology.

## Introduction

The development of digital pathology leads to accumulation of large-scale whole-slide imaging data, laying the foundation of big data for computational pathology. Rich morphological features buried in whole-slide image (WSI) provide diagnostic information of the disease and guidance on the decision for treatment. Advances in deep learning algorithms enable the analyses of gigapixel WSIs at scale for disease diagnosis^1,2^, prognosis^3–6^ and treatment selection^7,8^.

Deep learning approaches have achieved human-level performance in recognizing natural images on the ImageNet competition^9–12^. However, automatic recognition of WSI remains challenging due to the super-high spatial resolution of WSI as compared with images from ImageNet^9^. To address this challenge, researchers divided WSI into small image patches and subsequently aggregated features of image patches to obtain slide-level feature.^4,13–16^ For example, Campanella and colleagues used standard multiple-instance learning (MIL) for diagnosis of prostate cancer, basal cell carcinoma and auxiliary lymph node metastasis of breast cancer by first ranking image patches with regards to slide-level labels and using the most relevant image patch for slide-level classification.^1^ Lu and colleagues developed a data-efficient weakly supervised approach^17^ for slide-level classification by attention-based pooling^18^ of all image patches instead of the most relevant patch used by standard MIL^1^. Based on this approach, Lu and colleagues developed TOAD for predicting tissue-of-origins for cancer of unknown primary.^19^ Meanwhile, this attention-based MIL method has been utilized for addressing the diagnostic tasks for cardiac allograft rejection screening in WSIs^7^ and prognostic prediction by fusing WSIs with different modalities of genomic data^3^. Apart from these diagnostic endeavors, analyses of large-scale WSIs have been proved to be feasible for the prediction of genetic markers. Coudray and colleagues reported a deep-learning-based approach for predicting somatic mutations in canonical driver genes for lung cancer via averaging the probabilities of image patches or counting the percentage of image patches classified as positive.^14^ In addition, multiple studies reported that micro-satellite instability can be predicted from WSIs in gastrointestinal cancer^20^, colorectal cancer^21–23^ and endometrial carcinoma^24^.

The transformer architecture designed for natural language understanding can capture long-range dependencies among different entities^25^. Transformer-based language architectures achieved superior performance in multiple language understanding tasks^25,26^. The self-attention operation is the key module underlying the success of transformer in that it captures dependencies in the input^25^. Although it was proposed for language understanding, transformer is task-agnostic. It has been widely adopted or revised for image recognition. Vision Transformer (ViT) is the direct adoption of transformer for image classification by splitting image into multiple patches and taking the flatten image patches as input^25^. Thereafter, ViT-based architectures have been widely used in medical imaging analyses.^27–29^

Here, we present an approach called <u>W</u>SI <u>I</u>nspection via <u>T</u>ransformer (WIT) for slide-level classification via holistically modeling dependencies among patches on the WSI. WIT takes as input the features of image patches that were extracted with an image model pretrained on ImageNet^30^. We collected a total number of 22,457 WSIs from TCGA, CPTAC and PANDA projects to develop and systematically evaluate WIT for detection of 32 cancer types and diagnosis of cancer. The TCGA consists of 11,623 WSIs covering 32 cancer types. The CPTAC dataset includes 3,414 WSIs from cancer patients and 1,638 WSIs from non-cancer controls. The PANDA dataset consists of 5,782 needle biopsy slides, 2891 of them are prostate cancers and rest are non-cancer controls. WIT achieved an accuracy of 82.1% in the detection of 32 cancer types on the TCGA dataset, 91.8% in diagnosis of cancer on the CPTAC dataset and 88.2% on the PANDA dataset, outperforming the attention-based MIL baseline by 31.6%, 5.4% and 9.3%, respectively. WIT can pinpoint the WSI regions that are most influential for its decision. WIT represents a new paradigm for computational pathology. It will facilitate the development of assistive tools for digital pathology.

## Results

### An overview of WIT

The procedures to develop WIT includes WSI segmentation and tiling, model development and evaluation (**Figure 1**). Firstly, we segmented WSI to identify tissue regions and subsequently tiled WSI into patches of 256 × 256 pixels (**Figure 1A**). WIT takes the flattened image patches as input. We used a pretrained model to extract a feature of 1024 dimensions for each image patch (See **Methods**). Meanwhile, the position embeddings of image patches on that WSI along with their extracted feature were fed into a transformer block. The transformer block consists of a multi-headed self-attention module and point-wise feed forward neural network. Residual connection is employed around these two sub-modules, followed by layer normalization^25^ (**Figure 1B**). The multi-headed self-attention module learns the dependencies among different image patches and the influence of each patch on the output such as slide labels (**Figure 1B**). WIT was evaluated for its capacity in slide classification and localization of image patches that exhibit significant association with slide labels (**Figure 1C**).

**Figure 1.**
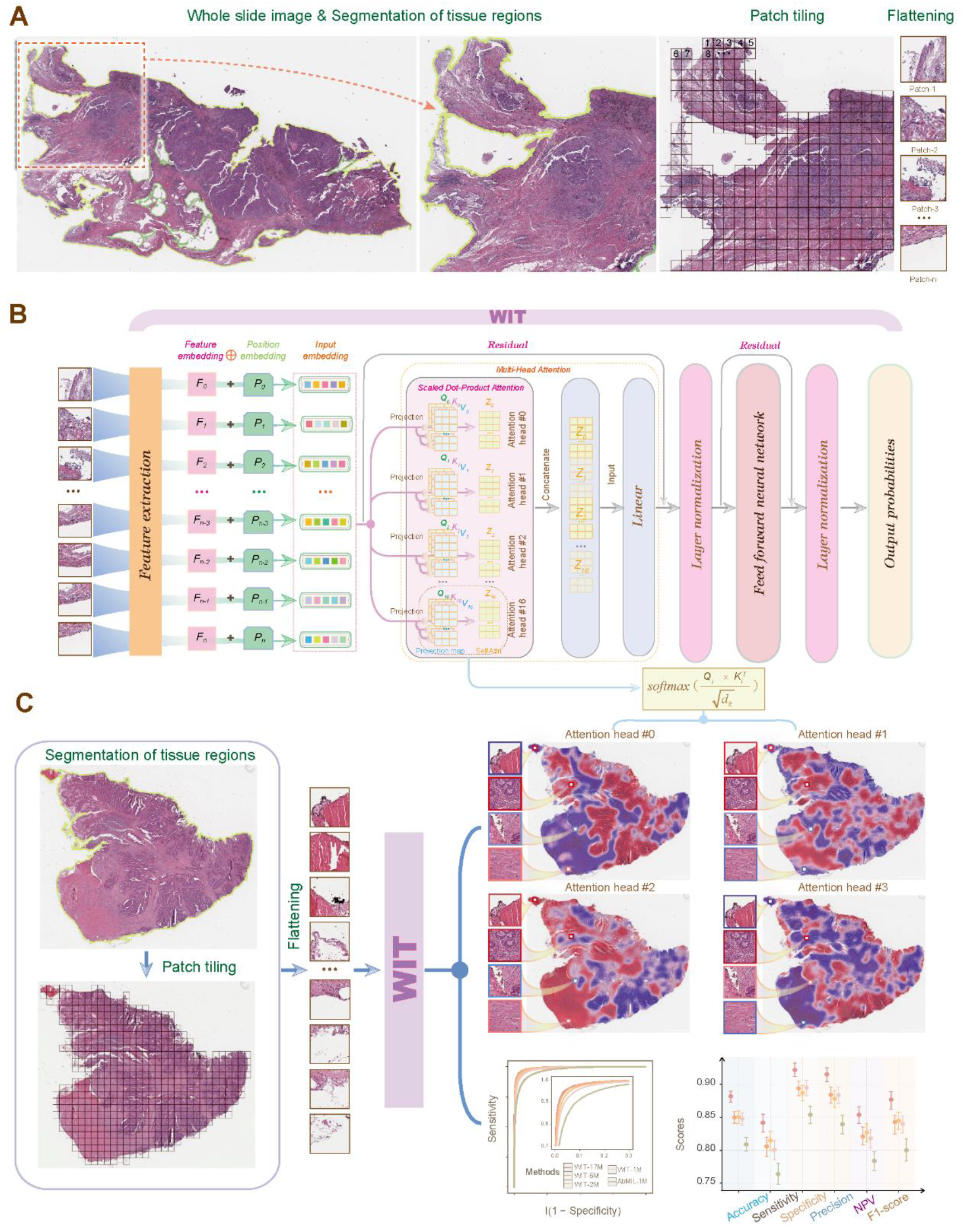
A flowchart illustrating the framework of WIT. (**A**) Illustration of the preprocessing steps: segmentation of tissue regions, patch tiling and flattening. (**B**) The architecture of WIT. (**C**) Evaluation of WIT for classification and model interpretability. WSI, Whole Slide Image; AbMIL, Attention-based Multiple Instance Learning.

### High performance of WIT in tissue-of-origin localization

We systematically evaluated the classification performance of WIT on The Cancer Genome Atlas (TCGA) dataset for tissue-of-origin localization via five-fold cross-validation (See **Methods**). The TCGA dataset consists of 11,623 formalin-fixed paraffin-embedded WSIs from 9565 individuals covering 32 cancer types (**Supplementary Table 1**). We examined classification performance of WIT with varying parameters such as 1, 2, 5 and 17 megabytes (**Supplementary Table 2**). We used the attention-based MIL model as the baseline model for comparison. The baseline has model parameters of 1 megabyte.

The accuracy of WIT was increasing with model size. Its top-1 accuracy was ranged from 73.1% (95% CI, 72.5% - 73.9%) for WIT-1Mb to 82.1% (80.7% - 83.3%) for WIT-17Mb while the baseline has a top-1 accuracy of 64.2% (60.6% - 66.0%) (**Figure 2A**). Top-2 and top-3 accuracies exhibited the same trend as top-1 accuracy (**Figure 2A and Supplementary Table 3**). Meanwhile, the micro-average AUROC of four WIT models are also higher than the baseline model (**Figure 2B**). WIT-17Mb achieved high performance in localization of 32 cancer types with respect to precision and recall rate (**Figure 2C**). WIT-17M achieved an average precision of 77.3% and recall rate of 75.6%, outperforming the baseline method by 29.5% and 37.5%, respectively. The confusion matrix of the baseline method was shown in **Supplementary Figure 1D**. WIT of different model size also had higher performance as compared with the baseline method when stratified by cancer types (**Figure 2D** and **Supplementary Table 4-6**). In addition, the F1 scores achieved by different WIT models are higher than the baseline method (**Figure 2E** and **Supplementary Table 7**). For example, WIT-1M had an average F1 score of 0.618 versus 0.554 as obtained by the baseline method, albeit WIT-1M and the baseline method had comparable model size.

**Figure 2.**
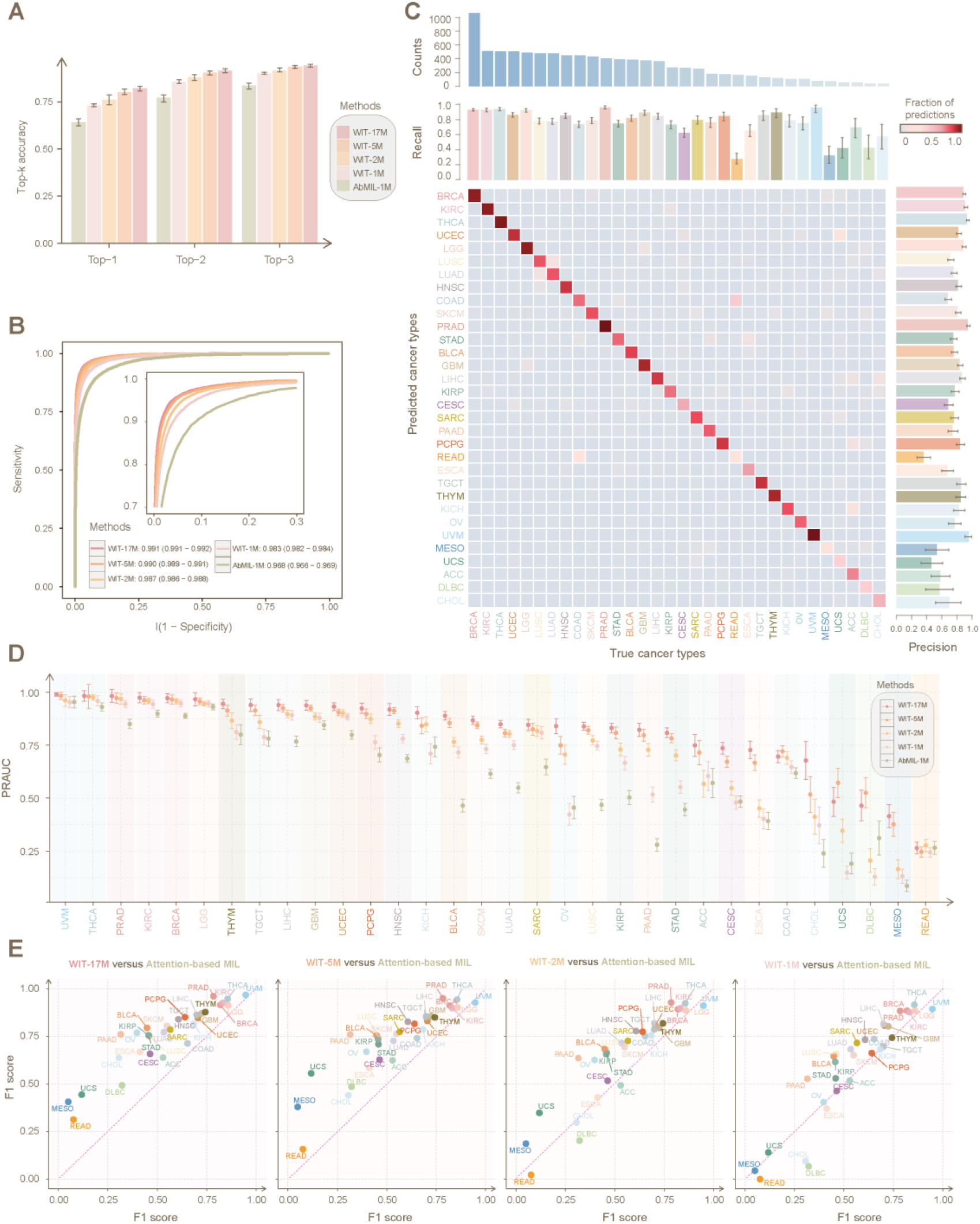
The classification performance of WIT in localization of tissue origins for 32 cancer types on TCGA dataset. (**A**) Top-*K* accuracy for localization of tumor origins, *K* ∈ {1, 2, 3}. (**B**) Micro-average area under the receiver operating curve. (**C**) Patient-level performance from five-fold cross-validation. Per origin count, precision and recall rate are plotted next the confusion matrix. The columns represent the true origin of the tumor and rows represent the prediction by the WIT model. (**D**) Area Under the Precision-Recall Curve (PRAUC) stratified by cancer types. (**D**) Scatter plots of F1 scores between different models. AbMIL, Attention-based Multiple Instance Learning.

### High performance of WIT in cancer diagnosis

WIT achieved high classification performance in the diagnosis of cancer on the CPTAC and PANDA datasets (See **Methods**). The CPTAC dataset consists of 5052 formalin-fixed paraffin-embedded WSIs from 1330 individuals (**Supplementary Table 8**). The PANDA dataset consists of 5,782 prostate WSIs subjected to needle biopsies^31^.

On the CPTAC dataset, WIT models achieved AUROCs ranged from 0.941 (95%CI, 0.934 – 0.949) to 0.953 (0.946 – 0.960) whereas the baseline model achieved an AUROC of 0.931 (0.931 - 0.969) (**Figure 3A**). WIT-17Mb achieved an accuracy of 0.918 (0.910 - 0.925) as compared with WIT models of smaller sizes as well as the baseline model. Similar trends were observed with respect to other classification metrics (**Figure 3B, C** and **Supplementary Table 9**). On the PANDA dataset, WIT-17Mb achieved the significantly higher AUROC as compared with WIT of smaller sizes the baseline model (DeLong’s test, all adjusted p-values < 2.2e-16, **Figure 3D**). Classification metrics such as accuracy, sensitivity, specificity, precision, negative predictive value and F1 score achieved by WIT-17Mb are also significantly higher than the other models (**Figure 3E, F** and **Supplementary Table 10**).

**Figure 3.**
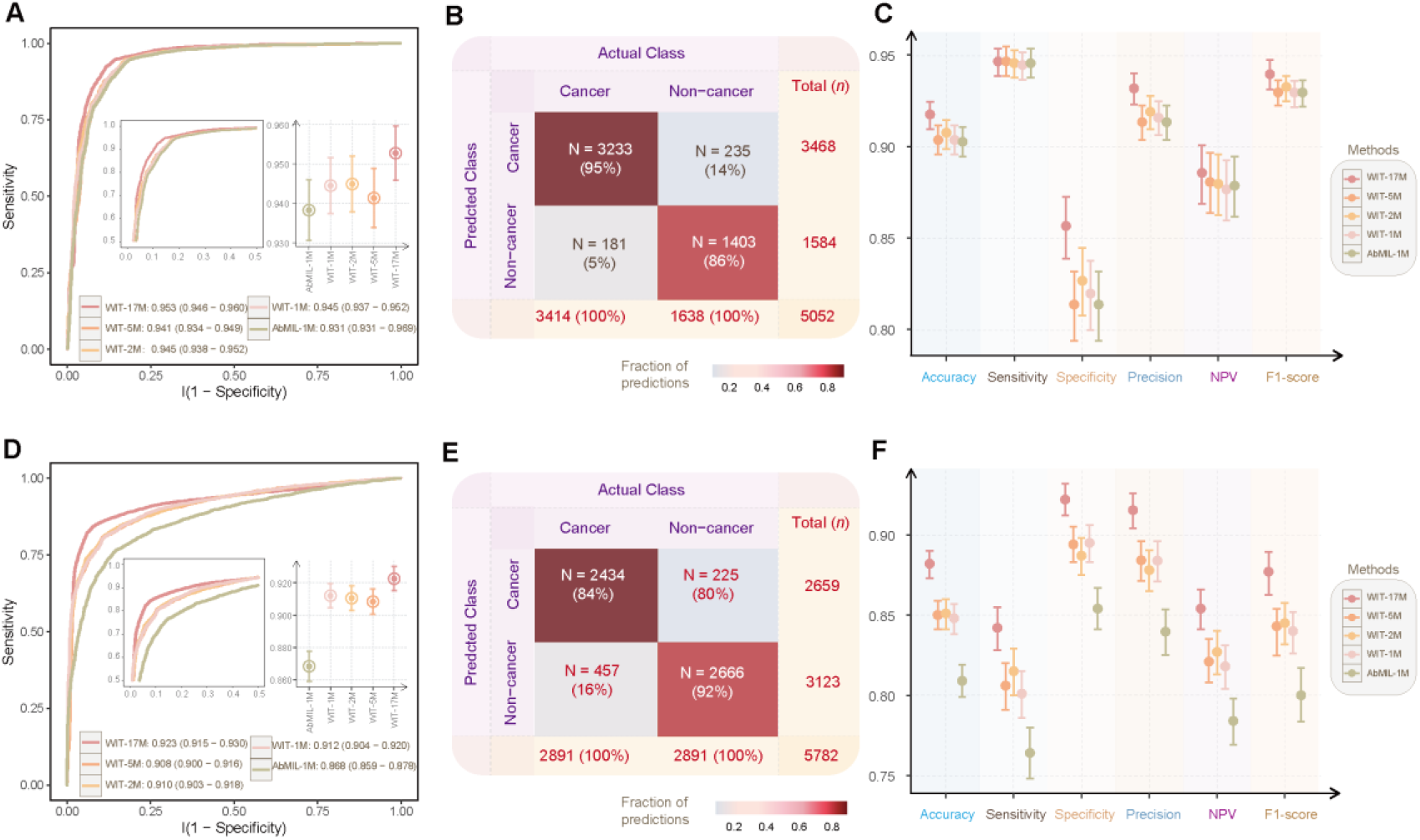
The classification performance of WIT in the diagnosis of cancer on CPTAC and PANDA datasets. (**A**, **D**) The receiver operating curves and area under the curves. (**B**, **E**) Confusion matrices. (**C, F**) Classification metrics of accuracy, sensitivity, specificity, precision, negative predictive value (NPV) and F1-score. AbMIL, Attention-based Multiple Instance Learning.

### Model interpretability

The multi-headed self-attention modules in WIT can quantify the importance of every image patch on the its prediction. We converted attention scores derived from WIT into human-interpretable heatmaps that highlights importance of WSI regions for prediction (See **Method**). In localization of 32 cancer types, WIT captures tumor regions that are considered to be morphology of different cancer types by pathologists in lung adenocarcinoma (LUAD, **Figure 4A**), rectum adenocarcinoma (**Figure 4B**), pancreatic adenocarcinoma (**Figure 4C**) and uterine corpus endometrial carcinoma (**Figure 4D**). For example, WIT identifies micropapillary tufts forming florets structure as strong evidence in detection of LUAD (**Figure 4A**). In the diagnosis of cancer, WIT pinpoints the tumor regions of non-keratinizing squamous cells with solid pattern in lung squamous cell carcinoma (**Figure 4E**) and confluent glandular and cribriform structure in UCEC (**Figure 5F**). In addition, WIT is able to identify prostate adenocarcinoma (**Figure 5G-H**) and a cluster of small poorly-formed glands (**Figure 5G**) from needle biopsy. We provided visualization of attention maps for a number of slides for exploration purpose in our interactive website (https://deeplearningplus.github.io/WIT-attention-maps/).

**Figure 4.**
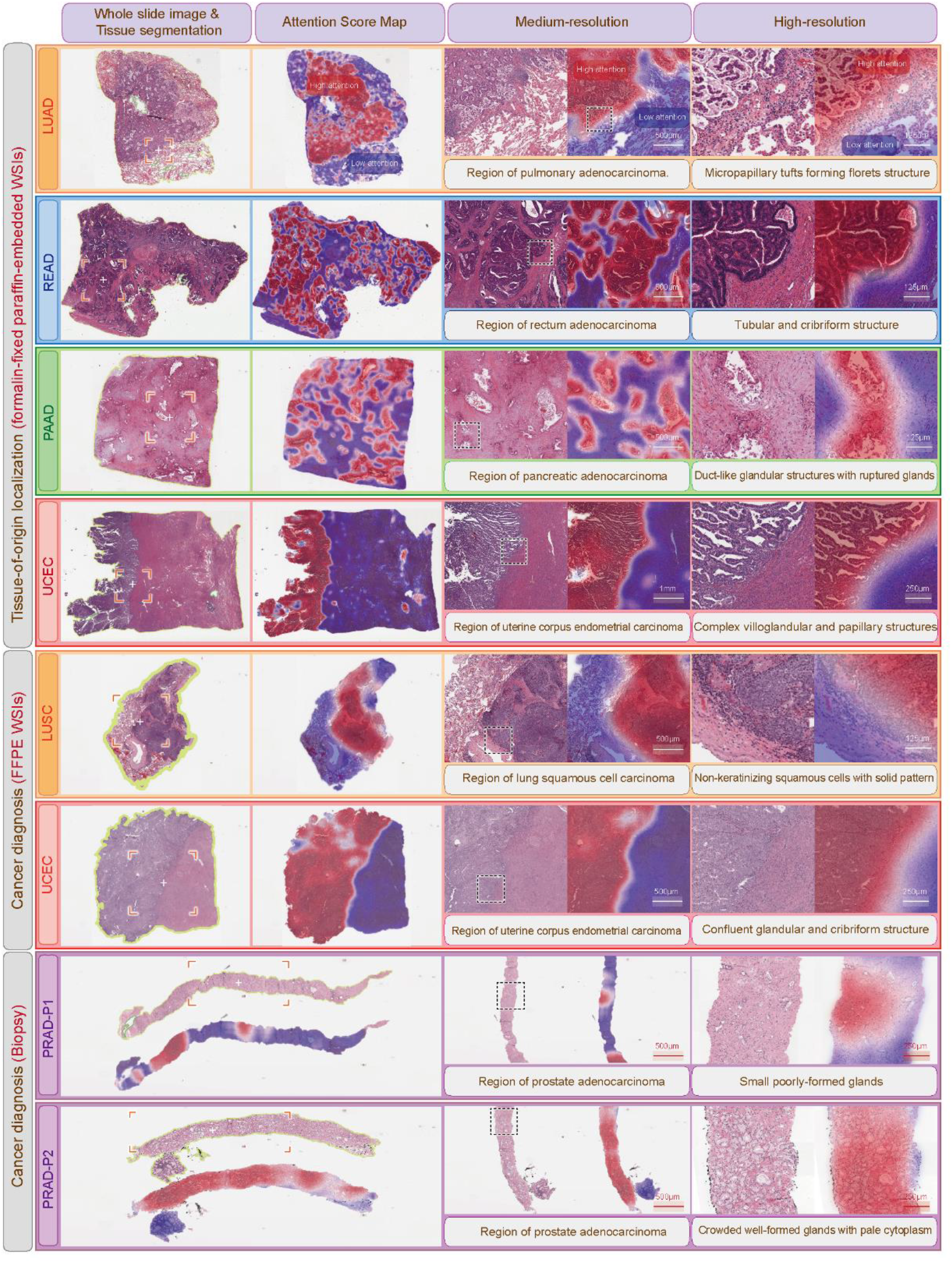
Attention maps of WIT for interpretability in localization of tissue origins and diagnosis of cancer from FFPE WSIs and biopsy. Boxes highlight the typical morphologic features corresponding to the textual description. The interactive visualization is available at https://deeplearningplus.github.io/WIT-attention-maps/.

## Discussion

In our study, we proposed a context-aware deep-learning approach WIT for slide-level localization of tumor origins and diagnosis of cancer from WSIs. WIT outperformed the attention-based MIL^19^ baseline by significant marginals across all classification tasks evaluated, especially in the detection of 32 cancer types where WIT achieved a micro-average AUROC of 0.991 (0.991-0.992) versus 0.968 (0.966 – 0.969) as obtained by the baseline method.

The high performance of WIT can be attributed to its context-aware ability to learn the potential nonlinear associations among image patches whereas the baseline method treats different image patches as independent instances. As WIT was built upon transformer^32^, the multi-headed self-attention module in transformer enables WIT to learn interrelation of patches in different subspaces while attention-based MIL is designed to aggregate multiple instances independently. Attention-based MIL methods have been widely and successfully adopted in addressing the challenges of computational pathology such as CLAM^17^, TOAD^19^ and CRANE^7^. WIT has the advantage of CLAM and TOAD in that it uses only the slide-level labels without any manual annotation. However, both CLAM and TOAD share the common limitations of MIL-based approaches^33^ in that they are context-independent but not context-aware.

WIT has several specific advantages. First, WIT can be easily scaled into models of different sizes. Large model has better classification performance as compared with smaller ones. However, the high performance of different WIT models cannot be merely attributed to their model sizes as compared with the attention-based MIL baseline. For example, in the detection of 32 cancer types, WIT-1Mb achieved significantly higher overall accuracy in comparison to the baseline method [73.1% (95% CI, 72.5%-73.9%) versus 64.2% (62.4%-66.0%)] although their model sizes are comparable. Therefore, the high performance of WIT is likely due to its ability to take into account nonlinear associations among all image patches. Besides, overall accuracy is steadily increasing with model size (**Supplementary Table 2**). Second, WIT is data-efficient in that we extracted image patches at ×20 magnification instead of full magnification. In this scenario, the 16 terabytes of TCGA WSI dataset were converted into a dataset of 200 gigabytes, enabling fast experimentation. Third, multi-head attentions used by WIT enable model interpretability from different feature representation subspaces, allowing for different morphological features to be identified by different attention heads. For example, we observed that one attention head of WIT identified micropapillary tufts forming florets structure as strong evidence for lung adenocarcinoma (**Figure 4A**) while the other heads pay attention to different tissue structure such as normal pulmonary alveoli (**Supplementary Figure 2**).

However, WIT was not without limitations. We used the ResNet50 model^30^ pretrained on the ImageNet dataset as feature extractor for image patches of WSI. The ImageNet is a collection of natural scene images. Therefore, it is definitely suboptimal by using this pretrained ResNet50 model^30^ in characterizing image patches clipped from WSIs. This strategy was also adopted by CLAM, TOAD and CRANE. Pretraining the feature extractor on image patches of WSIs may have the potential to improve the performance of WIT and all MIL-based methods. However, this will drastically increase the computational resources. We will address this issue in our future study. In addition, the 2D spatial dependencies among image patches is lost as WIT accepts flattened patches as input. Addressing this drawback with multi-dimensional transformers such as axial attention^34^ will improve the performance of WIT.

Weakly supervised learning such as MIL-based approaches have been successfully applied in addressing the challenges of computational pathology. However, their limitations are apparent in that they treat instances independently. Here, we addressed this challenge by presenting WIT – a deep-learning method based on transformer for learning feature presentation of whole slide via by taking into account nonlinear associations among image patches. WIT will facilitate adoption of deep-learning based solution and enable knowledge discovery in computational pathology.

## Methods

### Whole-slide image (WSI) preprocessing

The slide image was segmented for the tissue regions using the CLAM Python package. We used ×20 magnification. We cropped the WSI into 256×256 patches within the segmented tissue regions and flattened them into an array. We extracted a feature of 1024 dimensions for these image patches from the second residual layer of pretrained ResNet50 model^11,30^ on ImageNet dataset. The extracted features of image patches from a WSI were saved to disk file.

### WIT architecture

WIT consists of an embedding layer and a transformer encoder followed by a softmax layer.

#### Embedding layer

This layer takes as input the elementwise summation of image patch features and position embeddings of the flattened image patches. We used the pretrained ResNet50 model^11,25^ as feature extractor for image patches.

#### The transformer encoder

The encoder has two components: a multi-headed self-attention and a position-wise feed-forward neural network.

The *i^th^* self-attention head is formulated as^25^:

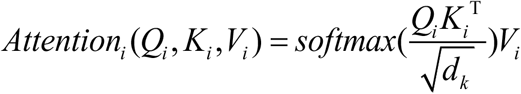

The input embeddings outputted from the embedding layer are projected to three matrices: query (*Q*_*i*_), key (*K*_*i*_) and value (*V*_*i*_). *d_k_* is the dimension of the query and it is used as scaling factor to mitigate the extreme small gradient^35^.

The multi-headed self-attention is the concatenation of multiple self-attention heads, allowing for the transformer attending to information in different feature representation subspaces. Multi-headed self-attention is formulated as^35^:

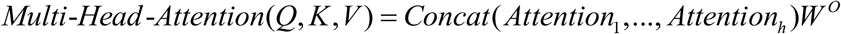

where 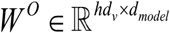 denotes the learned projection matrix.

The position-wise feed-forward neural network (*FFN*) consists of two linear layers with *ReLu* activation in-between:

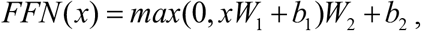

where *W_1_* and *W_2_* are weight matrices and *b*_1_ and *b*_2_ are the bias.

Layer-wise normalization^36^ is used in the front and rear of *FFN*. Residual connection^11^ is applied to improve information flow.

### Model training

The WSIs are random sampled and trained using WIT for 100 epochs. The weights and bias parameters of the model are initialized randomly, and the ground-truth label is slide-level labels. Cross-entropy loss^37^ is used in the classification task, the loss function captures how close the two distributions. The model parameters are updated via the *AdamW* optimizer with an initial learning rate of 2×10^−5^, weight decay of 1×10^−5^. WIT was trained with *PyTorch* (version 1.12.0) and *transformers* (version 4.21.1) on NVIDIA DGX A100.

### Different WIT Models

We evaluated four WIT models with different parameters by varying the hidden size: WIT-1Mb, WIT-2Mb, WIT-5Mb and WIT-17Mb. Details of these models are provided in **Supplementary Table 2**.

### WSI datasets

We collected a total number of 22,457 WSIs from The Cancer Genome Atlas (TCGA dataset, n = 11,623), The Clinical Proteomic Tumor Analysis Consortium (CPTAC dataset, n = 5,052) and PANDA (PANDA dataset, n = 5,782).

#### TCGA dataset

The TCGA dataset covers 32 cancer types: BRCA, KIRC, THCA, UCEC, LGG, LUSC, LUAD, HNSC, COAD, SKCM, PRAD, STAD, BLCA, GBM, LIHC, KIHC, CESC, SARC, PAAD, PCPG, READ, ESCA, TGCT, THYM, KICH, OV, UVM, MESO, UCS, ACC, DLBC and CHOL. The formalin-fixed paraffin embedded (FFPE) hematoxylin and eosin (H&E) stained WSIs are used. The details are in **Supplementary Table 1**.

#### CPTAC dataset

We collected a total of 11,623 WSIs from the Cancer Imaging Archive CPTAC Pathology Portal. The collected projects consisted of CPTAC-LUAD, CPTAC-LSCC, CPTAC-SAR, CPTAC-UCEC, CPTAC-UCEC, CPTAC-CCRCC, CPTAC-PDA, CPTAC-HNSCC, CPTAC-SAR and CPTAC-CM (**Supplementary Table 8**). The FFPE, H&E stained WSIs from normal donors and cancer patients are used.

#### The PANDA dataset

This dataset consists of 5,782 slides from prostate cancer patients and non-cancer individuals subjected to needle biopsies. There are 2,891 non-cancer biopsy WSIs. We randomly sampled 5,782 cancer biopsy WSIs to mitigate class imbalance cancer and non-cancer slides.

### Model evaluation

We used area under the receiver operating cureve (AUROC), accuracy, precision (also known as positive predictive value), recall rate, negative predictive value (NPV) and F1 score to assess the perfomance of WIT. Precision is the ratio of true positives to total predicted positives. Recall rate is the ratio of true positives to total actual positives. We reported the top-*K* accuracy for *K* = 1,2,3 on localization of 32 cancer types. NPV is defined as the number of true negatives divided by the number of samples predicted to be negative. F1-score is the harmonic mean of precision and recall rate.

### Baseline method

We used attention-based MIL^17–19^ implemented in TOAD^19^ as baseline method. Attention-based MILs are widely used in computational pathology studies. It takes a WSI as a bag and image patches on that WSI as instances. It uses attention-based pooling to aggregate the features of all image patches to obtain slide-level feature representations.

Let *H* = {*h*_1_,…, *h_k_* } be a bag of *K* instances, the MIL pooling is defined as^30^:

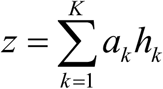

*a_k_* is the attention score for the *k^th^* instance, which is defined as ^30^:

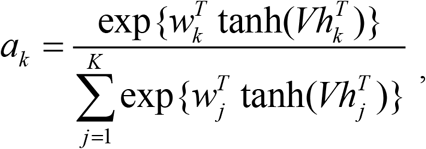

where ∀ *_k=1,…,K_*, *w*_K_ ∈ ℝ*_L×1_* and *V* ∈ ℝ*^L×M^* are parameters. The tanh is used as activation function. The network module is trained to assign an attention score *a_t_* for each patch ^30^:

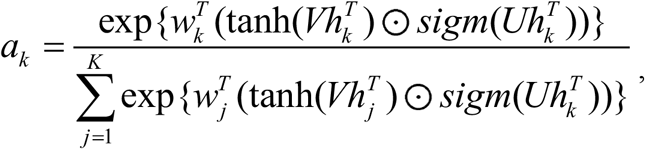

where *U* ∈ ℝ*^L×M^* are parameters, ⨀ is an element-wise multiplication and sigm(.) is sigmoid non-linearity.

### Visualization of attention map

For a given self-attention head, let *α* is the self-attention matrix; *α_i,j_* is the attention weight between the *i^th^* and *j^th^*. The attention score of the *i^th^* patch with slide-level representation *α_i,cls_* measures the contribution of the *i^th^* patch on classification. *CLS* stands for a slide-level representation where we added at the start of flattened feature array of image patches for each WSI, which is used for classification during training. The self-attention is obtained via:

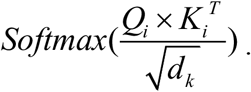

Assumed there are *K* patches in a WSI, the first row of each self-attention matrix (denoted as *α_0_*) quantifies the influence of each patch on classification. *α_0_* is converted to normalized percentile scores and scaled to the interval of [0,1] as proposed in CLAM^17^. The normalized attention scores were converted to RGB colours using a disperse colourmap values and displayed on the spatial regions in the slide with high attention displayed in red and low attention in deep purple using Matlibplot (version 3.5.2). We tiled the WSI into 256×256 patches using a overlap of 0.80 to create more fine-grained heatmaps. Gaussian blur is used to smooth uneven pixel values in a heatmap image using OpenSlide (version 3.4.1). The attention scores of heatmaps are claculated by NVIDIA DGX A100 with GPUs. We use the code of CLAM Python package for attention map visualization^17^.

### Statistical and software

We conducted our experiment with Python (version 3.8.10), OpenSlide (version 1.2.0), Pillow (version 9.1.1), R (version 4.2.1), ggplot2 (version 3.3.6), ROCR (version 1.0.11), multiROC (version 1.1.1) and PROC^38^ (version 1.18.0). The visualization of precision-recall curve (PRC) and calculation of area under PRC were performed with ROCR. Calculation of micro-averaged AUROC was performed with multiROC. Calculation of AUROC was performed with PROC.^38^ The 95% confidence intervals of the AUROC were calculated using DeLong’s methods implemented in pROC. The calculation of 95% confidence intervals for accuracy, sensitivity, specificity, precision, negative predictive value and F1 score with Clopper-Pearson method^39^.

## Acknowledgements

We are grateful for researchers for their generosity to made their data publicly available. This work was supported by the National Natural Science Foundation of China (Grant No. 32270688 and 31801117 to X.L. and 82073287 to Q.Z.), National Key Research and Development Program of China (Grant No. 2021YFC2500400 to K.C.), Program for Changjiang Scholars and Innovative Research Team in University in China (Grant No. IRT_14R40 to K.C.).

## Author Contributions

Xiangchun Li and Kexin Chen designed and supervised the study; Xiangchun Li and Hongru Shen performed data analysis and wrote the manuscript; Xiangchun Li developed the model; Jianghua Wu interpreted the whole-slide image data. Xiangchun Li, Hongru Shen, Xilin Shen, Jiani Hu, Jilei Liu and Qiang Zhang collected data; Yan Sun provided comments on the results. Hongru Shen, Xiangchun Li and Kexin Chen revised the manuscript.

## Data availability

The TCGA dataset is available at https://portal.gdc.cancer.gov. The CPTAC dataset is available at https://cancerimagingarchive.net/datascope/cptac. The PANDA dataset is available at https://www.kaggle.com/c/prostate-cancer-grade-assessment/data.

## Code availability

Code will be available at https://github.com/deeplearningplus/WIT.

## Figure legends

**Supplementary Figure 1. The classification performance of WIT in localization of tissue origins for 32 cancer types on TCGA dataset by WIT-5M (A), WIT-2M (B), WIT-1M (C) and Attention-based MIL (D)**.

**Supplementary Figure 2. Attention maps of WIT for interpretability in different attention heads.**

**Supplementary Table 1. The summary of patients in TCGA dataset.**

**Supplementary Table 2. The configurations of WIT with different model size.**

**Supplementary Table 3. Top-***K* **accuracy of WIT and AbMIL in tissue-of-origin localization.**

**Supplementary Table 4. Precision of WIT and AbMIL in in tissue-of-origin localization.**

**Supplementary Table 5. Recall rate of WIT and AbMIL in tissue-of-origin localization.**

**Supplementary Table 6. Area under the precision-recall curve for WIT and AbMIL in tissue-of-origin localization.**

**Supplementary Table 7. F1 score of WIT and AbMIL in tissue-of-origin localization.**

**Supplementary Table 8. The summary of patients in CPTAC dataset.**

**Supplementary Table 9. Classification metrics of WIT and AbMIL in cancer diagnosis on the CPTAC dataset.**

**Supplementary Table 10. Classification metrics of WIT and AbMIL in cancer diagnosis on the PANDA dataset**.

