## Supplementary information for "An efficient context-aware approach for whole slide image classification"

### Supplementary materials

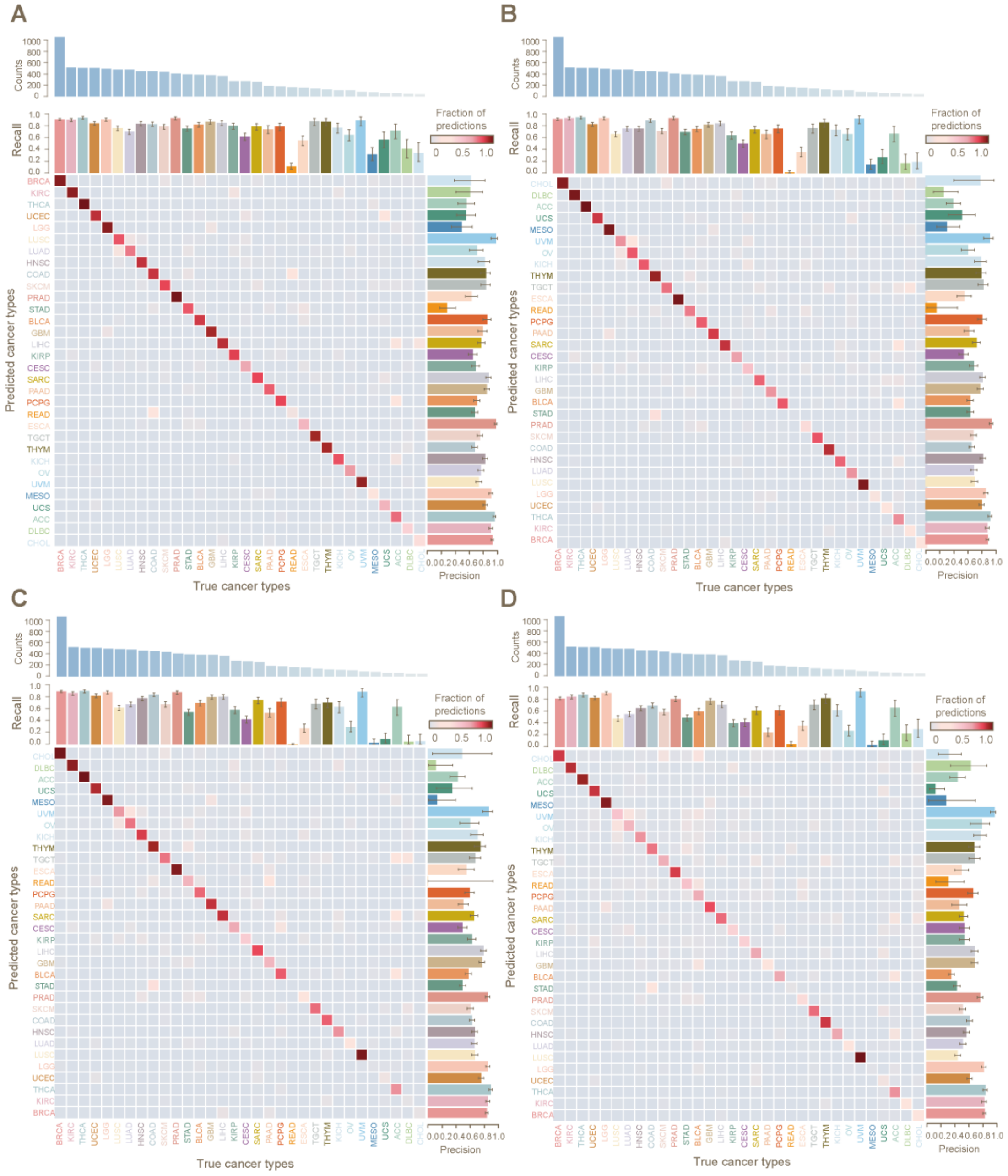


**Supplementary Figure 1. The classification performance of WIT in localization of tissue origins for 32 cancer types on TCGA dataset by WIT-5M (A), WIT-2M (B), WIT-1M (C) and Attention-based MIL (D).**

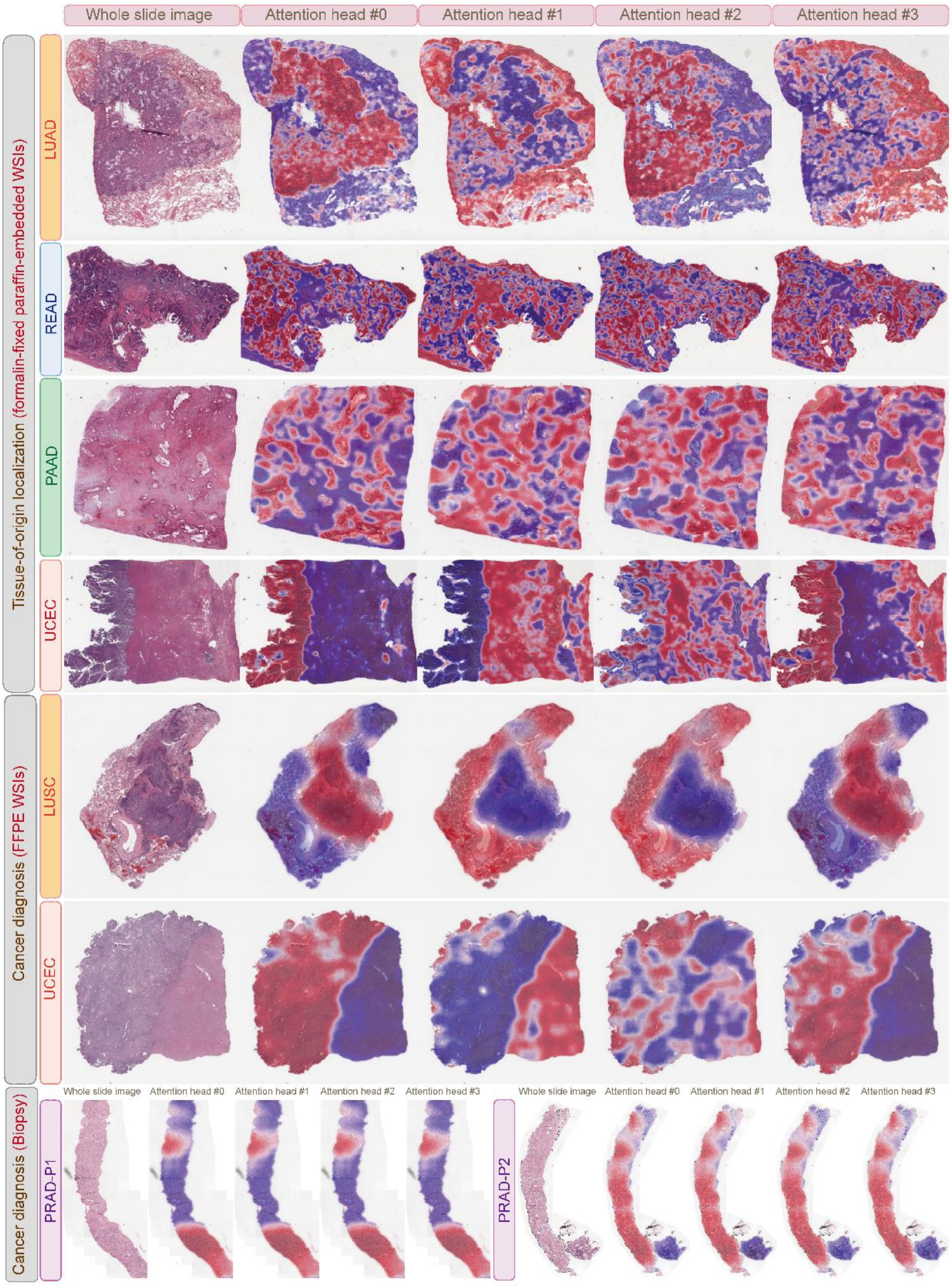


**Supplementary Figure 2. Attention maps of WIT for interpretability in different attention heads.**

**Supplementary Table 1. The summary of patients in TCGA dataset.**

| Abbreviation | Cancer types | Slides (no.) | Individual (no.) |
|---|---|---|---|

| | | | |
|---|---|---|---|
| ACC | Adrenocortical Carcinoma | 227 | 56 |
| BLCA | Bladder Urothelial Carcinoma | 455 | 384 |
| BRCA | Breast Invasive Carcinoma | 1131 | 1062 |
| CESC | Cervical Squamous Cell Carcinoma and Endocervical Adenocarcinoma | 279 | 269 |
| CHOL | Cholangiocarcinoma | 38 | 38 |
| COAD | Colon Adenocarcinoma | 456 | 448 |
| DLBC | Lymphoid Neoplasm Diffuse Large B-Cell Lymphoma | 42 | 42 |
| ESCA | Esophageal Carcinoma | 158 | 156 |
| GBM | Glioblastoma Multiforme | 817 | 378 |
| HNSC | Head and Neck Squamous Cell Carcinoma | 471 | 449 |
| KICH | Kidney Chromophobe | 121 | 109 |
| KIRC | Kidney Renal Clear Cell Carcinoma | 519 | 513 |
| KIRP | Kidney Renal Papillary Cell Carcinoma | 295 | 272 |
| LGG | Brain Lower Grade Glioma | 837 | 489 |
| LIHC | Liver Hepatocellular Carcinoma | 377 | 363 |
| LUAD | Lung Adenocarcinoma | 540 | 477 |
| LUSC | Lung Squamous Cell Carcinoma | 512 | 478 |
| MESO | Mesothelioma | 86 | 74 |
| OV | Ovarian Serous Cystadenocarcinoma | 107 | 106 |
| PAAD | Pancreatic Adenocarcinoma | 209 | 183 |
| PCPG | Pheochromocytoma and Paraganglioma | 196 | 176 |

| | | | |
|---|---|---|---|
| PRAD | Prostate Adenocarcinoma | 449 | 403 |
| READ | Rectum Adenocarcinoma | 164 | 163 |
| SARC | Sarcoma | 600 | 254 |
| SKCM | Skin Cutaneous Melanoma | 473 | 431 |
| STAD | Stomach Adenocarcinoma | 417 | 391 |
| TGCT | Testicular Germ Cell Tumors | 211 | 133 |
| THCA | Thyroid Carcinoma | 518 | 505 |
| THYM | Thymoma | 181 | 121 |
| UCEC | Uterine Corpus Endometrial Carcinoma | 566 | 505 |
| UCS | Uterine Carcinosarcoma | 91 | 57 |
| UVM | Uveal Melanoma | 80 | 80 |

**Supplementary Table 2. The configurations of WIT with different model size.**

| Model | Model name | Model size | Layer | Hidden | Embedding |
|---|---|---|---|---|---|
| WIT | WIT-17M | 16.80M | 1 | 1024 | 1024 |
| | WIT-5M | 4.99M | 1 | 512 | 1024 |
| | WIT-2M | 1.64M | 1 | 256 | 1024 |
| | WIT-1M | 1.07M | 1 | 192 | 1024 |

**Supplementary Table3. Top-*K* accuracy of WIT and AbMIL in tissue-of-origin localization.**

| Top-*K* accuracy | Method | Accuracy | Lower | Upper |
|---|---|---|---|---|
| Top-1 accuracy | WIT-17M | 0.821432 | 0.807109 | 0.832723 |
| | WIT-5M | 0.801777 | 0.789336 | 0.817041 |
| | WIT-2M | 0.76184 | 0.734971 | 0.787245 |
| | WIT-1M | 0.73058 | 0.724516 | 0.739153 |
| | AbMIL-1M | 0.642133 | 0.623628 | 0.659697 |

| | | | | |
|---|---|---|---|---|
| Top-2 accuracy | WIT-17M | 0.915944 | 0.905907 | 0.926294 |
| | WIT-5M | 0.905802 | 0.891793 | 0.913225 |
| | WIT-2M | 0.879352 | 0.863042 | 0.894929 |
| | WIT-1M | 0.855619 | 0.846315 | 0.868792 |
| | AbMIL-1M | 0.773027 | 0.749085 | 0.7862 |
| Top-3 accuracy | WIT-17M | 0.943544 | 0.934135 | 0.949294 |
| | WIT-5M | 0.936435 | 0.927339 | 0.943544 |
| | WIT-2M | 0.915944 | 0.910089 | 0.930476 |
| | WIT-1M | 0.900889 | 0.89702 | 0.907475 |
| | AbMIL-1M | 0.837637 | 0.817564 | 0.848406 |
| Top-4 accuracy | WIT-17M | 0.958913 | 0.953476 | 0.965499 |
| | WIT-5M | 0.951176 | 0.945635 | 0.957135 |
| | WIT-2M | 0.93999 | 0.932567 | 0.952431 |
| | WIT-1M | 0.924935 | 0.918975 | 0.930476 |
| | AbMIL-1M | 0.874334 | 0.861997 | 0.883429 |
| Top-5 accuracy | WIT-17M | 0.96644 | 0.961317 | 0.972295 |
| | WIT-5M | 0.96184 | 0.957135 | 0.966022 |
| | WIT-2M | 0.951908 | 0.945112 | 0.962886 |
| | WIT-1M | 0.940094 | 0.936749 | 0.943021 |
| | AbMIL-1M | 0.900889 | 0.896498 | 0.904861 |

**Supplementary Table 4. Precision of WIT and AbMIL in in tissue-of-origin localization.**

| | WIT-17M | WIT-5M | WIT-2M | WIT-1M | AbMIL-1M |
|---|---|---|---|---|---|
| BRCA | 0.904 (0.885 - 0.921) | 0.92 (0.902 - 0.936) | 0.884 (0.863 - 0.902) | 0.863 (0.842 - 0.883) | 0.841 (0.817 - 0.863) |
| KIRC | 0.924 (0.898 - 0.946) | 0.902 (0.873 - 0.926) | 0.879 (0.848 - 0.905) | 0.877 (0.846 - 0.904) | 0.835 (0.8 - 0.866) |

| | | | | | |
|---|---|---|---|---|---|
| THCA | 0.952 (0.929 - 0.969) | 0.955 (0.933 - 0.972) | 0.918 (0.891 - 0.94) | 0.92 (0.893 - 0.942) | 0.851 (0.817 - 0.881) |
| UCEC | 0.832 (0.797 - 0.863) | 0.825 (0.789 - 0.857) | 0.799 (0.762 - 0.833) | 0.781 (0.744 - 0.816) | 0.627 (0.588 - 0.664) |
| LGG | 0.902 (0.873 - 0.927) | 0.904 (0.874 - 0.928) | 0.86 (0.827 - 0.889) | 0.879 (0.847 - 0.906) | 0.833 (0.798 - 0.864) |
| LUSC | 0.724 (0.683 - 0.762) | 0.732 (0.691 - 0.771) | 0.699 (0.654 - 0.741) | 0.689 (0.643 - 0.732) | 0.453 (0.408 - 0.498) |
| LUAD | 0.769 (0.729 - 0.806) | 0.761 (0.719 - 0.801) | 0.689 (0.647 - 0.729) | 0.679 (0.635 - 0.721) | 0.528 (0.482 - 0.573) |
| HNSC | 0.83 (0.793 - 0.864) | 0.822 (0.784 - 0.856) | 0.817 (0.776 - 0.853) | 0.685 (0.643 - 0.725) | 0.579 (0.534 - 0.623) |
| COAD | 0.692 (0.648 - 0.733) | 0.673 (0.632 - 0.712) | 0.657 (0.617 - 0.695) | 0.648 (0.608 - 0.687) | 0.624 (0.58 - 0.667) |
| SKCM | 0.823 (0.783 - 0.859) | 0.746 (0.703 - 0.786) | 0.683 (0.638 - 0.726) | 0.621 (0.575 - 0.664) | 0.525 (0.479 - 0.571) |
| PRAD | 0.96 (0.936 - 0.977) | 0.976 (0.956 - 0.989) | 0.937 (0.908 - 0.959) | 0.878 (0.842 - 0.908) | 0.778 (0.735 - 0.817) |
| STAD | 0.762 (0.717 - 0.804) | 0.674 (0.627 - 0.717) | 0.637 (0.59 - 0.683) | 0.509 (0.461 - 0.558) | 0.44 (0.392 - 0.489) |
| BLCA | 0.772 (0.728 - 0.812) | 0.703 (0.659 - 0.745) | 0.637 (0.59 - 0.681) | 0.595 (0.549 - 0.641) | 0.363 (0.325 - 0.402) |
| GBM | 0.851 (0.812 - 0.885) | 0.845 (0.805 - 0.88) | 0.781 (0.736 - 0.821) | 0.792 (0.748 - 0.831) | 0.699 (0.651 - 0.743) |
| LIHC | 0.882 (0.844 - 0.914) | 0.872 (0.832 - 0.905) | 0.811 (0.768 - 0.85) | 0.818 (0.774 - 0.856) | 0.7 (0.65 - 0.746) |
| KIRP | 0.783 (0.728 - 0.833) | 0.686 (0.631 - 0.737) | 0.687 (0.625 - 0.744) | 0.644 (0.581 - 0.703) | 0.549 (0.476 - 0.621) |

| | | | | | |
|---|---|---|---|---|---|
| CESC | 0.697 (0.635 - 0.754) | 0.645 (0.583 - 0.703) | 0.543 (0.478 - 0.606) | 0.507 (0.44 - 0.573) | 0.548 (0.476 - 0.619) |
| SARC | 0.777 (0.721 - 0.826) | 0.765 (0.709 - 0.816) | 0.725 (0.666 - 0.779) | 0.677 (0.62 - 0.731) | 0.537 (0.477 - 0.596) |
| PAAD | 0.757 (0.688 - 0.817) | 0.785 (0.716 - 0.844) | 0.623 (0.55 - 0.692) | 0.516 (0.443 - 0.588) | 0.478 (0.371 - 0.586) |
| PCPG | 0.856 (0.795 - 0.905) | 0.852 (0.788 - 0.903) | 0.809 (0.74 - 0.866) | 0.61 (0.54 - 0.676) | 0.677 (0.598 - 0.749) |
| READ | 0.363 (0.278 - 0.454) | 0.277 (0.173 - 0.402) | 0.154 (0.019 - 0.454) | 0 (0 - 0.95) | 0.318 (0.139 - 0.549) |
| ESCA | 0.685 (0.603 - 0.758) | 0.63 (0.542 - 0.711) | 0.55 (0.447 - 0.65) | 0.566 (0.447 - 0.679) | 0.509 (0.41 - 0.608) |
| TGCT | 0.87 (0.8 - 0.923) | 0.835 (0.762 - 0.892) | 0.826 (0.747 - 0.889) | 0.694 (0.609 - 0.771) | 0.699 (0.614 - 0.776) |
| THYM | 0.864 (0.791 - 0.919) | 0.833 (0.757 - 0.894) | 0.797 (0.717 - 0.863) | 0.77 (0.681 - 0.844) | 0.693 (0.609 - 0.768) |
| KICH | 0.843 (0.758 - 0.908) | 0.814 (0.724 - 0.884) | 0.79 (0.697 - 0.865) | 0.729 (0.629 - 0.815) | 0.786 (0.683 - 0.868) |
| OV | 0.784 (0.692 - 0.86) | 0.701 (0.6 - 0.79) | 0.605 (0.509 - 0.696) | 0.615 (0.47 - 0.747) | 0.8 (0.631 - 0.916) |
| UVM | 0.975 (0.912 - 0.997) | 0.973 (0.905 - 0.997) | 0.912 (0.828 - 0.964) | 0.889 (0.8 - 0.948) | 0.986 (0.927 - 1) |
| MESO | 0.545 (0.388 - 0.696) | 0.489 (0.341 - 0.639) | 0.303 (0.156 - 0.487) | 0.133 (0.017 - 0.405) | 0.286 (0.037 - 0.71) |
| UCS | 0.471 (0.329 - 0.615) | 0.552 (0.415 - 0.683) | 0.517 (0.325 - 0.706) | 0.357 (0.128 - 0.649) | 0.133 (0.051 - 0.268) |
| ACC | 0.591 (0.463 - 0.71) | 0.556 (0.434 - 0.673) | 0.394 (0.294 - 0.5) | 0.434 (0.325 - 0.547) | 0.45 (0.338 - 0.565) |

| | | | | | |
|---|---|---|---|---|---|
| DLBC | 0.581 (0.391 - 0.755) | 0.607 (0.406 - 0.785) | 0.259 (0.111 - 0.463) | 0.118 (0.015 - 0.364) | 0.643 (0.351 - 0.872) |
| CHOL | 0.71 (0.52 - 0.858) | 0.619 (0.384 - 0.819) | 0.778 (0.4 - 0.972) | 0.5 (0.068 - 0.932) | 0.324 (0.174 - 0.505) |

**Supplementary Table 5. Recall rate of WIT and AbMIL in tissue-of-origin localization.**

| | WIT-17M | WIT-5M | WIT-2M | WIT-1M | AbMIL-1M |
|---|---|---|---|---|---|
| BRCA | 0.928 (0.911 - 0.943) | 0.905 (0.886 - 0.922) | 0.903 (0.884 - 0.92) | 0.91 (0.891 - 0.926) | 0.798 (0.773 - 0.822) |
| KIRC | 0.928 (0.902 - 0.949) | 0.897 (0.867 - 0.922) | 0.918 (0.891 - 0.94) | 0.877 (0.846 - 0.904) | 0.828 (0.793 - 0.86) |
| THCA | 0.943 (0.919 - 0.961) | 0.933 (0.907 - 0.953) | 0.931 (0.905 - 0.951) | 0.913 (0.885 - 0.936) | 0.861 (0.828 - 0.89) |
| UCEC | 0.863 (0.83 - 0.892) | 0.84 (0.805 - 0.871) | 0.818 (0.781 - 0.851) | 0.836 (0.8 - 0.867) | 0.808 (0.771 - 0.841) |
| LGG | 0.924 (0.897 - 0.946) | 0.902 (0.872 - 0.927) | 0.916 (0.888 - 0.939) | 0.892 (0.861 - 0.918) | 0.888 (0.856 - 0.914) |
| LUSC | 0.78 (0.74 - 0.817) | 0.755 (0.714 - 0.793) | 0.655 (0.61 - 0.697) | 0.63 (0.585 - 0.673) | 0.469 (0.423 - 0.514) |
| LUAD | 0.776 (0.736 - 0.812) | 0.696 (0.653 - 0.737) | 0.744 (0.703 - 0.783) | 0.688 (0.644 - 0.729) | 0.541 (0.495 - 0.586) |
| HNSC | 0.851 (0.814 - 0.882) | 0.833 (0.795 - 0.866) | 0.744 (0.701 - 0.784) | 0.788 (0.748 - 0.825) | 0.639 (0.593 - 0.684) |
| COAD | 0.737 (0.693 - 0.777) | 0.826 (0.788 - 0.86) | 0.879 (0.846 - 0.908) | 0.855 (0.819 - 0.886) | 0.685 (0.64 - 0.728) |
| SKCM | 0.789 (0.747 - 0.826) | 0.784 (0.742 - 0.822) | 0.705 (0.66 - 0.748) | 0.687 (0.641 - 0.73) | 0.575 (0.527 - 0.623) |

| | | | | | |
|---|---|---|---|---|---|
| PRAD | 0.963 (0.939 - 0.979) | 0.926 (0.895 - 0.949) | 0.921 (0.89 - 0.945) | 0.891 (0.856 - 0.92) | 0.792 (0.749 - 0.83) |
| STAD | 0.747 (0.701 - 0.789) | 0.749 (0.703 - 0.792) | 0.688 (0.639 - 0.734) | 0.552 (0.502 - 0.602) | 0.478 (0.428 - 0.529) |
| BLCA | 0.82 (0.778 - 0.857) | 0.815 (0.773 - 0.853) | 0.74 (0.693 - 0.783) | 0.708 (0.66 - 0.753) | 0.586 (0.535 - 0.636) |
| GBM | 0.892 (0.856 - 0.921) | 0.868 (0.829 - 0.9) | 0.81 (0.766 - 0.848) | 0.815 (0.772 - 0.853) | 0.754 (0.707 - 0.797) |
| LIHC | 0.846 (0.804 - 0.881) | 0.843 (0.801 - 0.879) | 0.829 (0.786 - 0.866) | 0.815 (0.772 - 0.854) | 0.7 (0.65 - 0.746) |
| KIRP | 0.732 (0.675 - 0.783) | 0.794 (0.741 - 0.841) | 0.629 (0.568 - 0.686) | 0.592 (0.531 - 0.651) | 0.39 (0.331 - 0.45) |
| CESC | 0.625 (0.564 - 0.683) | 0.613 (0.552 - 0.672) | 0.494 (0.433 - 0.556) | 0.428 (0.368 - 0.489) | 0.401 (0.342 - 0.463) |
| SARC | 0.795 (0.74 - 0.843) | 0.783 (0.728 - 0.833) | 0.728 (0.669 - 0.782) | 0.76 (0.702 - 0.811) | 0.598 (0.535 - 0.659) |
| PAAD | 0.765 (0.697 - 0.824) | 0.738 (0.668 - 0.8) | 0.65 (0.576 - 0.719) | 0.541 (0.466 - 0.615) | 0.235 (0.176 - 0.303) |
| PCPG | 0.847 (0.785 - 0.896) | 0.784 (0.716 - 0.842) | 0.744 (0.673 - 0.807) | 0.727 (0.655 - 0.792) | 0.608 (0.532 - 0.681) |
| READ | 0.276 (0.209 - 0.351) | 0.11 (0.067 - 0.169) | 0.012 (0.001 - 0.044) | 0 (0 - 0.018) | 0.043 (0.017 - 0.086) |
| ESCA | 0.654 (0.574 - 0.728) | 0.545 (0.463 - 0.625) | 0.353 (0.278 - 0.433) | 0.276 (0.207 - 0.353) | 0.346 (0.272 - 0.426) |
| TGCT | 0.857 (0.786 - 0.912) | 0.872 (0.803 - 0.924) | 0.752 (0.67 - 0.823) | 0.699 (0.614 - 0.776) | 0.699 (0.614 - 0.776) |
| THYM | 0.893 (0.823 - 0.942) | 0.868 (0.794 - 0.922) | 0.843 (0.766 - 0.903) | 0.719 (0.63 - 0.797) | 0.802 (0.719 - 0.869) |

| | | | | | |
|---|---|---|---|---|---|
| KICH | 0.789 (0.7 - 0.861) | 0.761 (0.67 - 0.838) | 0.725 (0.631 - 0.806) | 0.642 (0.545 - 0.732) | 0.606 (0.507 - 0.698) |
| OV | 0.755 (0.662 - 0.833) | 0.642 (0.543 - 0.732) | 0.651 (0.552 - 0.741) | 0.302 (0.217 - 0.399) | 0.264 (0.183 - 0.359) |
| UVM | 0.963 (0.894 - 0.992) | 0.887 (0.797 - 0.947) | 0.912 (0.828 - 0.964) | 0.9 (0.812 - 0.956) | 0.912 (0.828 - 0.964) |
| MESO | 0.324 (0.22 - 0.443) | 0.311 (0.208 - 0.429) | 0.135 (0.067 - 0.235) | 0.027 (0.003 - 0.094) | 0.027 (0.003 - 0.094) |
| UCS | 0.421 (0.291 - 0.559) | 0.561 (0.424 - 0.693) | 0.263 (0.155 - 0.397) | 0.088 (0.029 - 0.193) | 0.105 (0.04 - 0.215) |
| ACC | 0.696 (0.559 - 0.812) | 0.714 (0.578 - 0.827) | 0.661 (0.522 - 0.782) | 0.643 (0.504 - 0.766) | 0.643 (0.504 - 0.766) |
| DLBC | 0.429 (0.277 - 0.59) | 0.405 (0.256 - 0.567) | 0.167 (0.07 - 0.314) | 0.048 (0.006 - 0.162) | 0.214 (0.103 - 0.368) |
| CHOL | 0.579 (0.408 - 0.737) | 0.342 (0.196 - 0.514) | 0.184 (0.077 - 0.343) | 0.053 (0.006 - 0.177) | 0.289 (0.154 - 0.459) |

**Supplementary Table 6. Area under the precision-recall curve for WIT and AbMIL in tissue-of-origin localization.**

| Method | Cancer types | PRAUC | Lower limit | Upper limit |
|---|---|---|---|---|
| WIT-17M | BRCA | 0.972 | 0.951 | 0.994 |
| | KIRC | 0.975 | 0.950 | 0.999 |
| | THCA | 0.983 | 0.958 | 1.008 |
| | UCEC | 0.932 | 0.913 | 0.952 |
| | LGG | 0.967 | 0.936 | 0.999 |
| | LUSC | 0.839 | 0.822 | 0.857 |
| | LUAD | 0.849 | 0.831 | 0.867 |
| | HNSC | 0.919 | 0.891 | 0.947 |
| | COAD | 0.697 | 0.668 | 0.725 |

| | | | | |
|---|---|---|---|---|
| | SKCM | 0.868 | 0.849 | 0.887 |
| | PRAD | 0.983 | 0.952 | 1.014 |
| | STAD | 0.809 | 0.785 | 0.832 |
| | BLCA | 0.889 | 0.869 | 0.910 |
| | GBM | 0.938 | 0.915 | 0.961 |
| | LIHC | 0.940 | 0.915 | 0.965 |
| | KIRP | 0.832 | 0.808 | 0.855 |
| | CESC | 0.736 | 0.709 | 0.764 |
| | SARC | 0.846 | 0.819 | 0.873 |
| | PAAD | 0.823 | 0.791 | 0.854 |
| | PCPG | 0.925 | 0.900 | 0.949 |
| | READ | 0.264 | 0.236 | 0.291 |
| | ESCA | 0.728 | 0.686 | 0.769 |
| | TGCT | 0.940 | 0.911 | 0.970 |
| | THYM | 0.946 | 0.924 | 0.968 |
| | KICH | 0.904 | 0.875 | 0.933 |
| | OV | 0.840 | 0.805 | 0.875 |
| | UVM | 0.991 | 0.982 | 0.999 |
| | MESO | 0.415 | 0.360 | 0.470 |
| | UCS | 0.482 | 0.414 | 0.551 |
| | ACC | 0.749 | 0.698 | 0.801 |
| | DLBC | 0.464 | 0.391 | 0.537 |
| | CHOL | 0.678 | 0.588 | 0.768 |
| | BRCA | 0.967 | 0.946 | 0.987 |
| | KIRC | 0.963 | 0.946 | 0.981 |
| | THCA | 0.981 | 0.923 | 1.038 |
| WIT-5M | UCEC | 0.906 | 0.890 | 0.922 |
| | LGG | 0.957 | 0.942 | 0.973 |
| | LUSC | 0.821 | 0.802 | 0.839 |

| | | | | |
|---|---|---|---|---|
| | LUAD | 0.807 | 0.790 | 0.825 |
| | HNSC | 0.913 | 0.899 | 0.928 |
| | COAD | 0.721 | 0.695 | 0.747 |
| | SKCM | 0.844 | 0.828 | 0.859 |
| | PRAD | 0.973 | 0.949 | 0.998 |
| | STAD | 0.783 | 0.761 | 0.804 |
| | BLCA | 0.855 | 0.834 | 0.876 |
| | GBM | 0.925 | 0.911 | 0.939 |
| | LIHC | 0.925 | 0.906 | 0.944 |
| | KIRP | 0.809 | 0.781 | 0.836 |
| | CESC | 0.672 | 0.645 | 0.699 |
| | SARC | 0.826 | 0.798 | 0.854 |
| | PAAD | 0.797 | 0.770 | 0.825 |
| | PCPG | 0.899 | 0.877 | 0.920 |
| | READ | 0.246 | 0.222 | 0.270 |
| | ESCA | 0.667 | 0.630 | 0.705 |
| | TGCT | 0.914 | 0.889 | 0.940 |
| | THYM | 0.914 | 0.888 | 0.940 |
| | KICH | 0.842 | 0.804 | 0.880 |
| | OV | 0.750 | 0.708 | 0.792 |
| | UVM | 0.983 | 0.965 | 1.001 |
| | MESO | 0.376 | 0.320 | 0.433 |
| | UCS | 0.572 | 0.503 | 0.642 |
| | ACC | 0.715 | 0.658 | 0.773 |
| | DLBC | 0.525 | 0.454 | 0.597 |
| | CHOL | 0.517 | 0.433 | 0.602 |
| | BRCA | 0.959 | 0.946 | 0.972 |
| WIT-2M | KIRC | 0.960 | 0.947 | 0.972 |
| | THCA | 0.976 | 0.964 | 0.989 |

| | | | | |
|---|---|---|---|---|
| | UCEC | 0.901 | 0.884 | 0.917 |
| | LGG | 0.946 | 0.936 | 0.956 |
| | LUSC | 0.773 | 0.751 | 0.794 |
| | LUAD | 0.803 | 0.787 | 0.819 |
| | HNSC | 0.852 | 0.834 | 0.870 |
| | COAD | 0.690 | 0.661 | 0.719 |
| | SKCM | 0.775 | 0.756 | 0.794 |
| | PRAD | 0.969 | 0.944 | 0.993 |
| | STAD | 0.701 | 0.677 | 0.726 |
| | BLCA | 0.766 | 0.745 | 0.787 |
| | GBM | 0.886 | 0.869 | 0.902 |
| | LIHC | 0.898 | 0.880 | 0.916 |
| | KIRP | 0.730 | 0.701 | 0.759 |
| | CESC | 0.547 | 0.514 | 0.581 |
| | SARC | 0.818 | 0.792 | 0.843 |
| | PAAD | 0.729 | 0.698 | 0.759 |
| | PCPG | 0.875 | 0.853 | 0.897 |
| | READ | 0.276 | 0.248 | 0.304 |
| | ESCA | 0.451 | 0.410 | 0.492 |
| | TGCT | 0.860 | 0.828 | 0.891 |
| | THYM | 0.866 | 0.828 | 0.903 |
| | KICH | 0.849 | 0.814 | 0.884 |
| | OV | 0.706 | 0.661 | 0.751 |
| | UVM | 0.962 | 0.934 | 0.990 |
| | MESO | 0.164 | 0.120 | 0.208 |
| | UCS | 0.346 | 0.293 | 0.400 |
| | ACC | 0.567 | 0.504 | 0.631 |
| | DLBC | 0.205 | 0.150 | 0.260 |
| | CHOL | 0.412 | 0.329 | 0.495 |

| | | | | |
|---|---|---|---|---|
| | BRCA | 0.948 | 0.933 | 0.964 |
| | KIRC | 0.944 | 0.924 | 0.963 |
| | THCA | 0.960 | 0.936 | 0.983 |
| | UCEC | 0.886 | 0.870 | 0.901 |
| | LGG | 0.946 | 0.930 | 0.962 |
| | LUSC | 0.747 | 0.727 | 0.767 |
| | LUAD | 0.751 | 0.730 | 0.771 |
| | HNSC | 0.781 | 0.761 | 0.800 |
| | COAD | 0.647 | 0.617 | 0.677 |
| | SKCM | 0.732 | 0.710 | 0.754 |
| | PRAD | 0.945 | 0.928 | 0.962 |
| | STAD | 0.553 | 0.522 | 0.583 |
| | BLCA | 0.716 | 0.691 | 0.742 |
| | GBM | 0.879 | 0.858 | 0.900 |
| WIT-1M | LIHC | 0.892 | 0.869 | 0.914 |
| | KIRP | 0.666 | 0.635 | 0.698 |
| | CESC | 0.479 | 0.447 | 0.511 |
| | SARC | 0.809 | 0.781 | 0.837 |
| | PAAD | 0.516 | 0.480 | 0.552 |
| | PCPG | 0.764 | 0.732 | 0.796 |
| | READ | 0.244 | 0.222 | 0.265 |
| | ESCA | 0.402 | 0.355 | 0.449 |
| | TGCT | 0.789 | 0.753 | 0.824 |
| | THYM | 0.812 | 0.778 | 0.846 |
| | KICH | 0.710 | 0.661 | 0.758 |
| | OV | 0.422 | 0.371 | 0.474 |
| | UVM | 0.953 | 0.926 | 0.980 |
| | MESO | 0.131 | 0.097 | 0.165 |
| | UCS | 0.148 | 0.117 | 0.178 |

| | | | | |
|---|---|---|---|---|
| AbMIL-1M | ACC | 0.605 | 0.536 | 0.673 |
| | DLBC | 0.127 | 0.089 | 0.166 |
| | CHOL | 0.371 | 0.284 | 0.459 |
| | BRCA | 0.888 | 0.876 | 0.900 |
| | KIRC | 0.899 | 0.883 | 0.916 |
| | THCA | 0.931 | 0.912 | 0.950 |
| | UCEC | 0.799 | 0.778 | 0.819 |
| | LGG | 0.932 | 0.918 | 0.946 |
| | LUSC | 0.468 | 0.442 | 0.494 |
| | LUAD | 0.549 | 0.525 | 0.573 |
| | HNSC | 0.687 | 0.668 | 0.706 |
| | COAD | 0.618 | 0.593 | 0.643 |
| | SKCM | 0.615 | 0.594 | 0.637 |
| | PRAD | 0.851 | 0.828 | 0.873 |
| | STAD | 0.445 | 0.416 | 0.474 |
| | BLCA | 0.465 | 0.434 | 0.495 |
| | GBM | 0.845 | 0.825 | 0.866 |
| | LIHC | 0.768 | 0.742 | 0.793 |
| | KIRP | 0.503 | 0.469 | 0.536 |
| | CESC | 0.483 | 0.457 | 0.509 |
| | SARC | 0.647 | 0.609 | 0.684 |
| | PAAD | 0.279 | 0.250 | 0.309 |
| | PCPG | 0.704 | 0.671 | 0.737 |
| | READ | 0.265 | 0.235 | 0.295 |
| | ESCA | 0.392 | 0.350 | 0.433 |
| | TGCT | 0.781 | 0.745 | 0.818 |
| | THYM | 0.800 | 0.753 | 0.848 |
| | KICH | 0.743 | 0.696 | 0.790 |
| | OV | 0.455 | 0.402 | 0.508 |

| | | | | |
|---|---|---|---|---|
| | UVM | 0.955 | 0.927 | 0.982 |
| | MESO | 0.085 | 0.058 | 0.111 |
| | UCS | 0.190 | 0.141 | 0.239 |
| | ACC | 0.572 | 0.502 | 0.641 |
| | DLBC | 0.311 | 0.230 | 0.392 |
| | CHOL | 0.238 | 0.173 | 0.304 |

**Supplementary Table 7. F1 score of WIT and AbMIL in tissue-of-origin localization.**

| | WIT-17M | WIT-5M | WIT-2M | WIT-1M | AbMIL-1M |
|---|---|---|---|---|---|
| BRCA | 0.916 | 0.913 | 0.893 | 0.886 | 0.819 |
| KIRC | 0.926 | 0.899 | 0.898 | 0.877 | 0.832 |
| THCA | 0.947 | 0.944 | 0.924 | 0.917 | 0.856 |
| UCEC | 0.847 | 0.832 | 0.808 | 0.808 | 0.706 |
| LGG | 0.913 | 0.903 | 0.887 | 0.885 | 0.859 |
| LUSC | 0.751 | 0.744 | 0.676 | 0.658 | 0.460 |
| LUAD | 0.772 | 0.727 | 0.716 | 0.683 | 0.534 |
| HNSC | 0.840 | 0.827 | 0.779 | 0.733 | 0.607 |
| COAD | 0.714 | 0.741 | 0.752 | 0.737 | 0.653 |
| SKCM | 0.806 | 0.765 | 0.694 | 0.652 | 0.549 |
| PRAD | 0.962 | 0.950 | 0.929 | 0.884 | 0.785 |
| STAD | 0.755 | 0.709 | 0.662 | 0.530 | 0.458 |
| BLCA | 0.795 | 0.755 | 0.684 | 0.647 | 0.448 |
| GBM | 0.871 | 0.856 | 0.795 | 0.803 | 0.725 |
| LIHC | 0.864 | 0.857 | 0.820 | 0.817 | 0.700 |
| KIRP | 0.757 | 0.736 | 0.656 | 0.617 | 0.456 |
| CESC | 0.659 | 0.629 | 0.518 | 0.464 | 0.464 |
| SARC | 0.786 | 0.774 | 0.727 | 0.716 | 0.566 |
| PAAD | 0.761 | 0.761 | 0.636 | 0.528 | 0.315 |

| | | | | | |
|---|---|---|---|---|---|
| PCPG | 0.851 | 0.817 | 0.775 | 0.663 | 0.641 |
| READ | 0.314 | 0.158 | 0.023 | - | 0.076 |
| ESCA | 0.669 | 0.584 | 0.430 | 0.371 | 0.412 |
| TGCT | 0.864 | 0.853 | 0.787 | 0.697 | 0.699 |
| THYM | 0.878 | 0.850 | 0.819 | 0.744 | 0.743 |
| KICH | 0.815 | 0.787 | 0.756 | 0.683 | 0.684 |
| OV | 0.769 | 0.670 | 0.627 | 0.405 | 0.397 |
| UVM | 0.969 | 0.928 | 0.913 | 0.894 | 0.948 |
| MESO | 0.407 | 0.380 | 0.187 | 0.045 | 0.049 |
| UCS | 0.444 | 0.557 | 0.349 | 0.141 | 0.118 |
| ACC | 0.639 | 0.625 | 0.493 | 0.518 | 0.529 |
| DLBC | 0.493 | 0.486 | 0.203 | 0.068 | 0.321 |
| CHOL | 0.638 | 0.441 | 0.298 | 0.095 | 0.306 |

**Supplementary Table 8. The summary of patients in CPTAC dataset.**

| Abbreviations | Cancer types | Individuals (no.) | Slides (no.) | Normal (no.) | Cancer (no.) |
|---|---|---|---|---|---|
| CCRCC | Clear Cell Renal Cell Carcinoma | 221 | 736 | 233 | 503 |
| CM | Cutaneous Melanoma | 93 | 378 | 95 | 283 |
| HNSCC | Head and Neck Squamous Cell Carcinoma | 107 | 334 | 97 | 237 |
| LSCC | Laryngeal Squamous Cell Carcinoma | 181 | 965 | 341 | 624 |
| LUAD | Lung Adenocarcinoma | 229 | 1041 | 371 | 670 |
| PDA | Pancreatic Ductal Adenocarcinoma | 164 | 487 | 150 | 337 |
| SAR | Sarcoma | 89 | 294 | 81 | 213 |
| UCEC | Endometrial Cancer | 246 | 817 | 270 | 547 |

**Supplementary Table 9. Classification metrics of WIT and AbMIL in cancer diagnosis on the CPTAC dataset.**

| Method | Classification metrics | Score | Lower limit | Upper limit |
|---|---|---|---|---|
| WIT-17M | accuracy | 0.918 | 0.910 | 0.925 |
| | sensitivity | 0.947 | 0.939 | 0.954 |
| | specificity | 0.857 | 0.839 | 0.873 |
| | precision | 0.932 | 0.923 | 0.940 |
| | NPV | 0.886 | 0.869 | 0.901 |
| | F1-score | 0.940 | 0.932 | 0.948 |
| WIT-5M | accuracy | 0.904 | 0.896 | 0.912 |
| | sensitivity | 0.947 | 0.939 | 0.955 |
| | specificity | 0.814 | 0.794 | 0.832 |
| | precision | 0.914 | 0.904 | 0.923 |
| | NPV | 0.881 | 0.864 | 0.897 |
| | F1-score | 0.930 | 0.923 | 0.937 |
| WIT-2M | accuracy | 0.908 | 0.899 | 0.915 |
| | sensitivity | 0.946 | 0.938 | 0.953 |
| | specificity | 0.827 | 0.808 | 0.845 |
| | precision | 0.919 | 0.910 | 0.928 |
| | NPV | 0.880 | 0.863 | 0.896 |
| | F1-score | 0.933 | 0.925 | 0.939 |
| WIT-1M | accuracy | 0.904 | 0.896 | 0.912 |
| | sensitivity | 0.945 | 0.937 | 0.952 |
| | specificity | 0.820 | 0.800 | 0.838 |
| | precision | 0.916 | 0.907 | 0.925 |
| | NPV | 0.877 | 0.860 | 0.893 |
| | F1-score | 0.930 | 0.922 | 0.936 |

| | | | | |
|---|---|---|---|---|
| AbMIL-1M | accuracy | 0.903 | 0.895 | 0.911 |
| | sensitivity | 0.946 | 0.938 | 0.954 |
| | specificity | 0.814 | 0.794 | 0.832 |
| | precision | 0.914 | 0.904 | 0.923 |
| | NPV | 0.879 | 0.862 | 0.895 |
| | F1-score | 0.930 | 0.922 | 0.937 |

**Supplementary Table 10. Classification metrics of WIT and AbMIL in cancer diagnosis on the PANDA dataset.**

| Method | Classification metrics | Score | Lower limit | Upper limit |
|---|---|---|---|---|
| WIT-17 | accuracy | 0.882 | 0.873 | 0.890 |
| | sensitivity | 0.842 | 0.828 | 0.855 |
| | specificity | 0.922 | 0.912 | 0.932 |
| | precision | 0.915 | 0.904 | 0.926 |
| | NPV | 0.854 | 0.841 | 0.866 |
| | F1-score | 0.877 | 0.863 | 0.889 |
| WIT-5M | accuracy | 0.850 | 0.841 | 0.859 |
| | sensitivity | 0.806 | 0.791 | 0.820 |
| | specificity | 0.894 | 0.883 | 0.905 |
| | precision | 0.884 | 0.871 | 0.896 |
| | NPV | 0.821 | 0.808 | 0.835 |
| | F1-score | 0.843 | 0.825 | 0.854 |
| WIT-2M | accuracy | 0.851 | 0.841 | 0.860 |
| | sensitivity | 0.815 | 0.800 | 0.829 |
| | specificity | 0.887 | 0.875 | 0.898 |
| | precision | 0.878 | 0.865 | 0.890 |
| | NPV | 0.827 | 0.813 | 0.840 |
| | F1-score | 0.845 | 0.832 | 0.858 |

| | | | | |
|---|---|---|---|---|
| WIT-1M | accuracy | 0.848 | 0.838 | 0.857 |
| | sensitivity | 0.801 | 0.786 | 0.815 |
| | specificity | 0.895 | 0.883 | 0.906 |
| | precision | 0.884 | 0.871 | 0.896 |
| | NPV | 0.818 | 0.804 | 0.831 |
| | F1-score | 0.840 | 0.826 | 0.852 |
| AbMIL-1M | accuracy | 0.809 | 0.799 | 0.819 |
| | sensitivity | 0.764 | 0.748 | 0.780 |
| | specificity | 0.854 | 0.841 | 0.867 |
| | precision | 0.840 | 0.825 | 0.853 |
| | NPV | 0.784 | 0.769 | 0.798 |
| | F1-score | 0.800 | 0.784 | 0.817 |
